# How The Transition of Aphid Life Cycles Affect Endosymbiotic Bacteria

**DOI:** 10.1101/2025.04.02.646943

**Authors:** Xin Tong, Shin-ichi Akimoto, Tomonari Nozaki, Masanori Arita, Shuji Shigenobu

## Abstract

How symbiotic bacteria coordinate and respond to their hosts during complex life cycle transitions is a fascinating and unresolved topic, particularly for organisms that exist solely in natural environments, where their intricate life cycles make laboratory cultivation extremely challenging. Complex life cycles are often associated with host alternation in aphids, especially in eriosomatine gall-forming aphids, which exhibit seven distinct morphs throughout their life cycle while shifting their host plants seasonally. Despite these dramatic changes, these aphids maintain the obligate symbiotic bacterium, *Buchnera aphidicola*, to enhance their nutritional status. The bacterial communities in our wild-derived samples were dominated by aphid symbionts, including *Buchnera, Regiella inseticola, Wolbachia, Arsenophonus*, and *Serratia symbiotica. Buchnera* and *Regiella* prevailed in *Tetraneura akinire* and *Tetraneura sorini*, while *Buchnera* and *Arsenophonus* were more abundant in *Eriosoma harunire*. Our study, utilizing 16S rRNA metagenomic sequencing on the aphid *Tetraneura akinire*, demonstrates that *Buchnera* plays significant roles during most life cycle stages but is absent during the sexual male stage, where it is replaced by *Regiella*. Moreover, *Buchnera* densities exhibit significant variability across the aphids’ life cycles and are notably lower in gall generations. This result contradicts the previous hypothesis that galls provide superior nutrition compared to aphids in grassroot-feeding generations. In this study, we reveal the dynamics of symbiotic bacteria across the complex life cycles of three eriosomatine aphid species that form galls on the same Japanese elm tree, *Ulmus davidiana* var. *japonica*. Within the framework of tripartite symbiosis, the seemingly passive symbiotic bacteria adjust the host’s specific life stages and influence the changing host ecology in a cascading manner.

## 1 INTRODUCTION

The beauty and complexity of symbiotic interactions in nature are well-documented due to their prevalence. These relationships enhance the survival and adaptability of the organisms involved and play crucial roles in individual development as well as shaping ecosystems. For example, some symbionts have a remarkable ability to expand the physiological capacity of their hosts, allowing them to exploit new metabolic and ecological niches (Stewart *et al*. 2005; Relman 2008). Correspondingly, many organisms with complex life cycles can capitalize on local resource utilization and an opportunity to exploit different ecological niches by shifting between different morphs in response to environmental changes (Forcada *et al*. 2008; Bell *et al*. 2014; Nettle & Bateson 2015;). For example, parasites with complex life cycles are common in nature, by exhibiting discrete ontogenetic changes and alternation between niches in response to environmental changes (Wilbur 1980; Moran 1992; Minelli & Fusco 2010; Auld & Tinsley 2015). The increased lifecycle complexity provides organisms with adaptive potentials to minimize negative environmental effects, including offering immediate benefits of dispersal ability after gaining mobile morphology, and long-term benefits with acquiring diverse genotypic chances for sexual reproduction (Rauch *et al*. 2005; Parker & Brisson 2019; Hanna & Abouheif 2021; Benesh *et al*. 2022). Such dramatic lifecycle transitions may entail huge cost and risk for increased selection pressure, leading to trade-offs (Moran 1994; Gandon 2004; Forcada *et al*. 2008; Benesh *et al*. 2022). Moreover, evolutionary adaptation in complex life cycles may be also constrained during lifecycle transition. For example, adaptative characters promoted in one life stage in a complex life cycle may compromise the evolution to other life stages as limits of ontogenetic reorganization adapting to various environmental changes (Bonett & Blair 2017; Fabre *et al*. 2004).

Aphid life cycles have evolved to be highly complex and diverse, including transitions between unique reproductive modes of cyclical parthenogenesis and viviparity, extreme polymorphism, eu/social behaviors, and obligate seasonal alternation between distinct host plant taxa (Hille Ris Lambers 1966; Moran 1992; Heie 1996; Jousselin *et al*. 2010; Favret 2022). Notably, many aphids are selective on host plant ranges and are often monophagous in one life stage but evolve to have two distinct sets of host plants such as shifting obligately between woody plants and grasses across life cycles (Akimoto 1985; Moran 1988; Stern *et al*. 2007; Aoki & Kurosu 2010). Alternating host plants seasonally was hypothesized to optimize nutrition through complementary host growth, but the cost of host alternation can also be very high, as about only 1 in 300 aphids succeed in returning to the primary host plants from the secondary host plants in early autumn (Ward et al, 1998). Moran (1988; 1992) suggested such obligate host-plant alternation was a consequence of seasonal polymorphism, as ancestral foundress morph may be constrained on the primary host plant while summer viviparous aphids acquire different host plants. More striking about aphids, some aphids can modify host plant by reprogramming its development and *de novo* inducing plant organs that are called galls (Stone & Schönrogge 2003; Harris & Pitzschke 2019; Korgaonkar *et al*. 2020). Gall formation represents unique lifecycle character and ability of aphids to dramatically modify host plant development. Aphid galls are not only species-specific in morphology but also tend to have different nutrient profiles by the inducing species even on the same host plant, though usually in a limited niche (Larson & Whitham 1991; Akimoto 1995; Suzuki *et al*. 2009). Galls induced by eriosomatine aphids on Japanese elm trees contain high concentrations in amino acids compared to unmodified leaves. The lifted amino-acid levels in galls may enhance aphid reproductive fitness as found in *Tetraneura* galling generations that produce larger numbers of offspring in highly nutritious galls (Koyama *et al*. 2004; Suzuki *et al*. 2009). Nonetheless, aphid gall formation and host alternation tend to occur together: almost all gall-forming aphids maintain high complexity in life cycle by seasonally shifting host plants and exhibiting extreme polymorphism, even with incredible ability to reprogram host plants (Moran 1988,1992;). Host plants may resist aphids from continuous feeding in galls (Williams and Whitham 1986; Goggin 2007). High cost of life-cycle transitions and species-specific modification on galls drive trade-offs the evolution of complex life cycle in aphids.

Although shifting across extreme polymorphism in complex life cycles, almost all aphids maintain endosymbiotic bacterial partners, the obligate nutritional symbiont *Buchnera aphidicola* starting from 200 million years ago, and some have other facultative symbionts such as *Serratia, Wolbachia* spp, *Arsenophonus* and *Regiella insectacola* (Douglas 1998; Douglas *et al*. 2006; Shigenobu *et al*. 2000; Manzano-Marin *et al*. 2017, 2023). Many of symbiotic bacteria in aphids are intracellular and can play essential roles through aphid life cycles (McLean *et al*. 2017). *Buchnera* promotes the nutritional supply in plant phloem sap especially by synthesizing essential amino acids and vitamins to support aphid growth (Douglas 1998, 2006; Douglas *et al*. 2001; Koga *et al*. 2003; Hansen & Moran 2011; Wilson *et al*. 2010). Despite facing the event of the reducing genomes, *Buchnera* maintains almost all genes related to biosynthesis of essential amino acids, and the pattern of gene loss is significantly nonrandom (Francis *et al*. 2001; Gil *et al*. 2002; Chong *et al*. 2019). Furthermore, *Buchnera* status is closely tied to aphid fitness. Concurrent decrease of aphid reproduction and *Buchnera* population in pea aphids, *Acyrthosiphon pisum*, after heat-shock treatment showed positive correlation between fitness of aphids and *Buchnera*, with more severe results for *Buchnera*. The survival rates of aphids and *Buchnera* population both fluctuated after being transferred to other host plants but became stable after a few generations (Zhang *et al*. 2016; Nousheen *et al*. 2022). How *Buchnera* responses to the evolution of life cycles across lineages remains yet to be determined.

Many facultative symbionts are essential to enhance aphid fitness as co-obligate partner to complement roles of *Buchnera* by biosynthesizing vitamin Bs and even improving survival against heat stress and parasite infection (Oliver *et al*. 2003; Vorburger *et al*. 2009; Manzano-Marin *et al*. 2017, 2023; Yorimoto *et al*. 2022). Co-infections of various bacterial symbionts can commonly exist in wild aphids (Liu *et al*. 2022), and the same is observed in lab-induced infection (McLean *et al*. 2017). To maximize fitness, however, aphid hosts may need to entail tight regulation on symbiont population in different developmental stages and environmental conditions. Moreover, microbiome assembly at the community level is highly predictable and convergent with similar patterns to primary succession observed in human infants, honeybees, and bumblebees (Stewart *et al*. 2018; Hammer *et al*. 2022), while stochastic colonization is evident at the strain level (Obadia *et al*. 2017; Vega & Gore 2017; Hammer *et al*. 2022). Vertical transmission of core symbionts predominates while horizontal transmission is evident through physical interaction with nestmates or environmental infection (Song *et al*. 2013; Kwong & Moran 2016), though the ratio of the core symbiont species shifts with aging process in worker bumblebees (Hammer *et al*. 2021). Different patterns of microbiome assemblies may cooperate hosts for colonization and persistence to corresponding life-cycle transitions.

To investigate how symbiotic bacteria coordinate and respond to their hosts during complex life cycle transitions, this study will focus on three gall-forming species that exhibit polymorphism and complex life cycles: *Tetraneura akinire, Tetraneura sorini* and *Eriosoma harunire*. We selected these three eriosomatine aphid species because they share the same primary host plant, the Japanese elm tree, where they form galls as their food source, but they utilize different secondary host plants in later life stage. Additionally, our familiarity with their natural habitats enables us to collect samples throughout their life cycle transitions in the field annually. Firstly, we performed community-level microbiome assays on the three gall-forming aphid species, to address gaps of symbiont diversity across the life cycles in wild-derived aphid samples. Secondly, we quantified the titers of the obligate symbiotic bacterium, *Buchnera*, throughout the aphid life cycle, focusing on three distinct categories of generations: the galling generations (foundresses and their offspring within the galls), the root generations (where aphids feed on the roots of certain grasses), and the sexual generations. Notably, *Tetraneura* aphids induce closed true galls, whereas *E. harunire* aphids generate ‘pseudo galls,’ including leaf-curl galls. According to Suzuki *et al*. (2009), these galls exhibit distinct amino acid profiles.

To understand how symbiotic microbes behave in response to complex frameworks of host life histories can be fundamental in evolutionary biology.

## 3 Results

### 3.1 Microbial community associated with *T.akinire, T.sorini* and *E.harunire* across life cycles

Despite the potential for diverse microbial interactions, the bacterial communities in our wild-derived samples remained relatively simple, dominated by known aphid symbionts, including *Buchnera, Regiella, Wolbachia, Arsenophonus*, and *Serratia* (Figure 1). Across all life stages, *Buchnera*, the maternally inherited primary symbiont, was the most abundant bacterium. However, in *T. akinire* males, *Regiella* became the dominant symbiont, while male *T. sorini* aphids exhibited an unexpectedly high diversity of bacterial taxa, including non-aphid-specific symbionts. Microbial composition also varied between aphids collected from tree galls and grass roots. In *T. akinire* and *T. sorini, Buchnera* and *Regiella* were the predominant bacteria, whereas in *E. harunire, Buchnera* and *Arsenophonus* were the most abundant.

To further evaluate symbiont diversity, we analyzed the V1-V2 and V3-V4 regions of the 16S rRNA gene separately. In the V1-V2 region, *Buchnera* remained the dominant symbiont, with relative abundance ranging from 32 to 2509 reads across different samples (Figure 1a). Other aphid-associated symbionts were detected at lower levels, including *Regiella, Wolbachia*, and *Arsenophonus*. Notably, in *T. sorini* males, *Regiella* became the dominant species (1458 reads), with a much lower abundance of *Buchnera* (32 reads). In *E. harunire* a large number of unassigned reads (2347) suggested either sequencing artifacts or the presence of unidentified microbes. In the V3-V4 region, *Buchnera* remained the most abundant, but we observed greater variability in other bacterial taxa (Figure 1b). *Regiella* showed a notable increase in *T. sorini*, with S10 harboring 3267 reads, while *Buchnera* was nearly absent. In *E. harunire, Buchnera* and *Arsenophonus* were the top symbionts, with *Buchnera* abundance ranging from 741 to 3102 reads. A high proportion of “Others” (3185 reads) in S5 suggests the presence of additional unidentified symbionts or environmental contamination.

**Fig 1.**
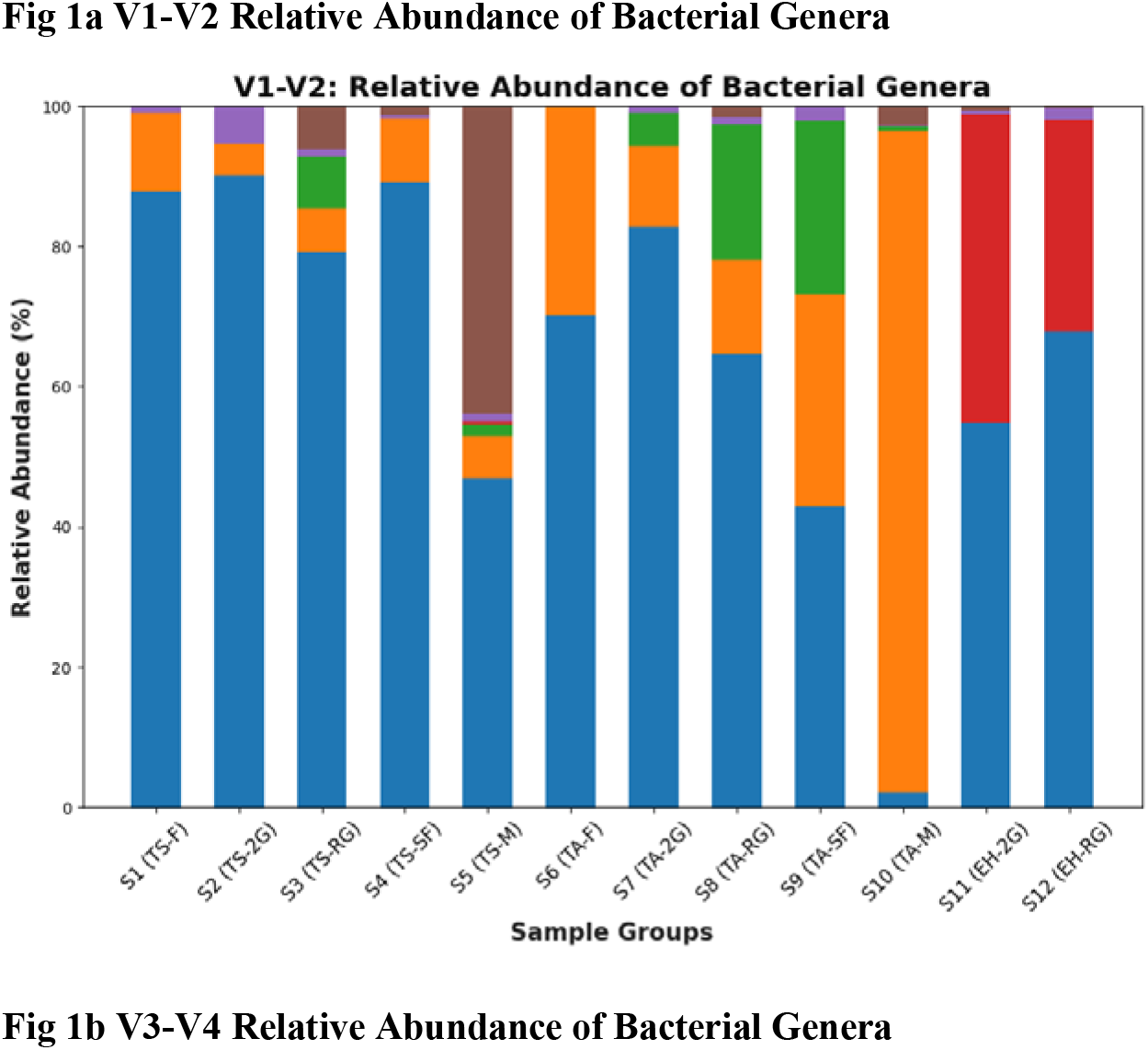
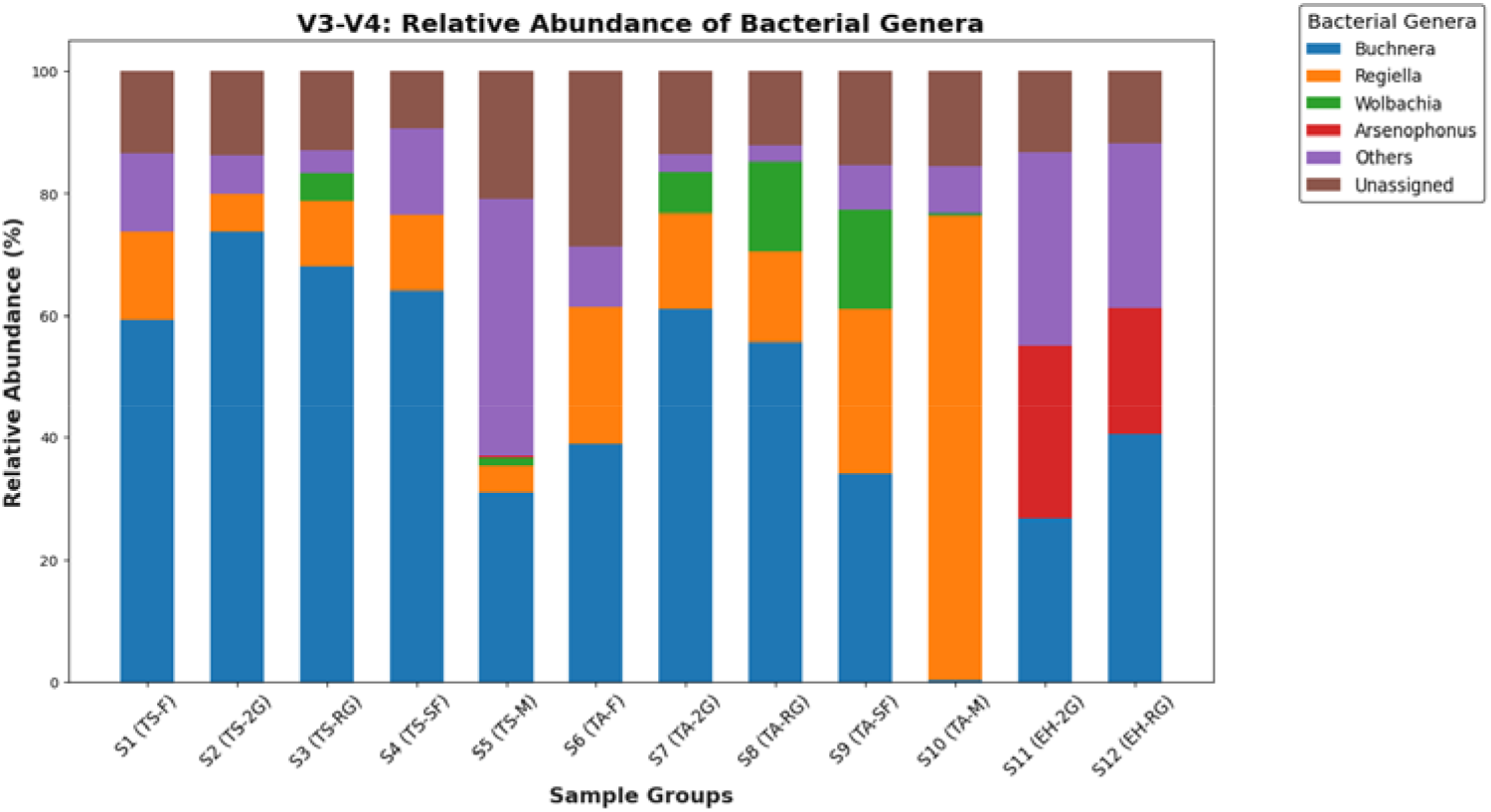
Taxonomic composition and relative abundance of symbiotic bacteria in *T. akinire, T. sorini*, and *E. harunire* across life stages by 16S rRNA metagenomic sequencing performed separately for the V1-V2 and V3-V4 regions. Five life stages were sampled for *Tetraneura* species, while only two life stages were sampled for *E. harunire* due to collection difficulties. See Table 1 for detailed sample information.

### 3.2 Diversity Analysis

We performed Kruskal-Wallis tests and pairwise comparisons to assess the differences in Shannon entropy values across species (TS, TA, EH), life stages (1^st^ generation, 2^nd^ generation, Root, Sexual_female, Male), and host groups (Elm gall, Grass root, No_feeding). The Kruskal-Wallis test indicated no significant difference in the diverssity between species (EH vs TA: 0.0528, EH vs TS: 0.2453, TA vs TS: 0.6015), life stages (e.g., 1st vs 2nd: 0.0833, 1st vs male: 0.1213, male vs sexual_female: 0.4386), or host groups (Elm gall vs Grass root: 0.8815, Elm gall vs No_feeding: 0.0500, Grass root vs No_feeding: 0.0771). Additionally, the q-values adjusted for multiple comparisons confirmed the lack of significant differences across these factors.

**Table 1.** Sample Collection Overview for *T. sorini, T. akinire*, and *E. harunire*. The samples listed below are mixed for 16S rRNA metagenomic sequencing, while the 4th instar and 4-day-old samples were used for qPCR quantification. ID represents the sample sequencing number, with samples of the same number pooled together for sequencing.

| ID | Species | Life stage | Instar | Site type | Collection Location | Collection Date |
| --- | --- | --- | --- | --- | --- | --- |
| S1 | *T. sorini* | Foundress | 4th instar | Elm leaf gall | Shiroishi, Sapporo / Iwamizawa | 2020.05.30~31 |
| S1 | *T. sorini* | Foundress | Adult | Elm leaf gall | Shiroishi, Sapporo / Iwamizawa | 2020.05.30~31 |
| S2 | *T. sorini* | 2nd generation | 4th instar | Elm leaf gall | Shiroishi, Sapporo | 2020.06.10 |
| S3 | *T. sorini* | Sexupara | 4th instar | *Miscanthus sinensis* root | Bibai City | 2020.10.02 |
| S3 | *T. sorini* | Apterous exule | Adult | *Miscanthus sinensis* root | Shiroishi, Sapporo | 2020.09.23 |
| S4 | *T. sorini* | Sexual female | 4-day-old | Elm tree bark | Mikasa City | 2020.10.14 |
| S5 | *T. sorini* | Male | 4-day-old | Elm tree bark | Mikasa City | 2020.10.14 |
| S6 | *T. akinire* | Foundress | 4th instar | Elm leaf gall | Hokkaido Univ, Sapporo | 2020.05.28 |
| S6 | *T. akinire* | Foundress | Adult | Elm leaf gall | Hokkaido Univ, Sapporo | 2020.05.28 |
| S7 | *T. akinire* | 2nd generation | 4th instar | Elm leaf gall | Hokkaido Univ, Sapporo | 2020.06.15 |
| S8 | *T. akinire* | Sexupara | 4th instar | *Setaria faberi* root | Hokkaido Univ, Sapporo | 2020.10.07 |
| S8 | *T. akinire* | Apterous exule | Adult | *Setaria faberi* root | Hokkaido Univ, Sapporo | 2020.10.07 |
| S9 | *T. akinire* | Sexual female | 4-day-old | Elm tree bark | Hokkaido Univ, Sapporo | 2020.10.14 |
| S10 | *T. akinire* | Male | 4-day-old | Elm tree bark | Hokkaido Univ, Sapporo | 2020.10.14 |
|  | *E.* |  |  |  |  |  |
| S11 | *harunire* | Foundress | Adult | Elm leaf gall | Hokkaido Univ, Sapporo | 2020.06.30 |
|  | *E.* |  |  |  |  |  |
| S11 | *harunire* | 2nd generation | 4th instar | Elm leaf gall | Hokkaido Univ, Sapporo | 2020.06.30 |
|  | *E.* |  |  |  |  |  |
| S12 | *harunire* | Sexupara | 4th instar | *Plantago asiatica* root | Hokkaido Univ, Sapporo | 2020.09.30 |
|  | *E.* |  |  |  |  |  |
| S12 | *harunire* | Apterous exule | Adult | *Plantago asiatica* root | Hokkaido Univ, Sapporo | 2020.09.15 |

A PERMANOVA test was conducted to assess the effects of species, life stage, and host on community composition across all samples. The analysis revealed a significant difference in community composition between species (pseudo-F = 8.50, p-value = 0.001), indicating that species contribute substantially to the variation observed in community structure (Fig. 2). In contrast, no significant differences were found for life stage (pseudo-F = 0.42, p-value = 0.999) or host (pseudo-F = 0.71, p-value = 0.685). Similar results were obtained for the *Tetranuera*-only dataset, where no significant differences in community composition were observed across life stages (pseudo-F = 0.36, p-value = 0.98) (Fig. 3).

**Figure 2.**
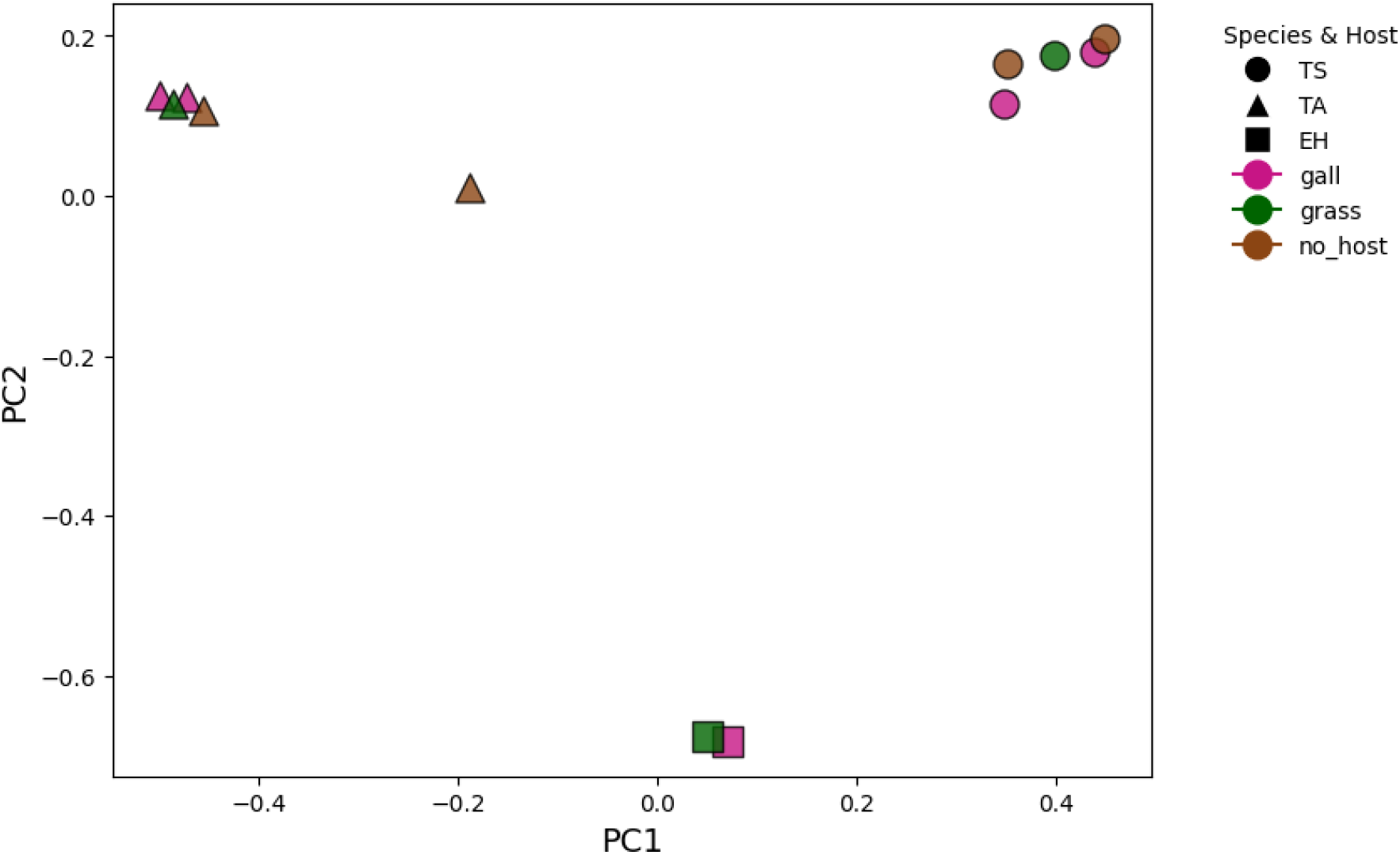
PCoA Analysis of Species and Host Influence on Community Composition. This figure presents the results of the Principal Coordinates Analysis (PCoA) for species and host across all samples, illustrating the variation in community composition based on these factors.

**Figure 3:**
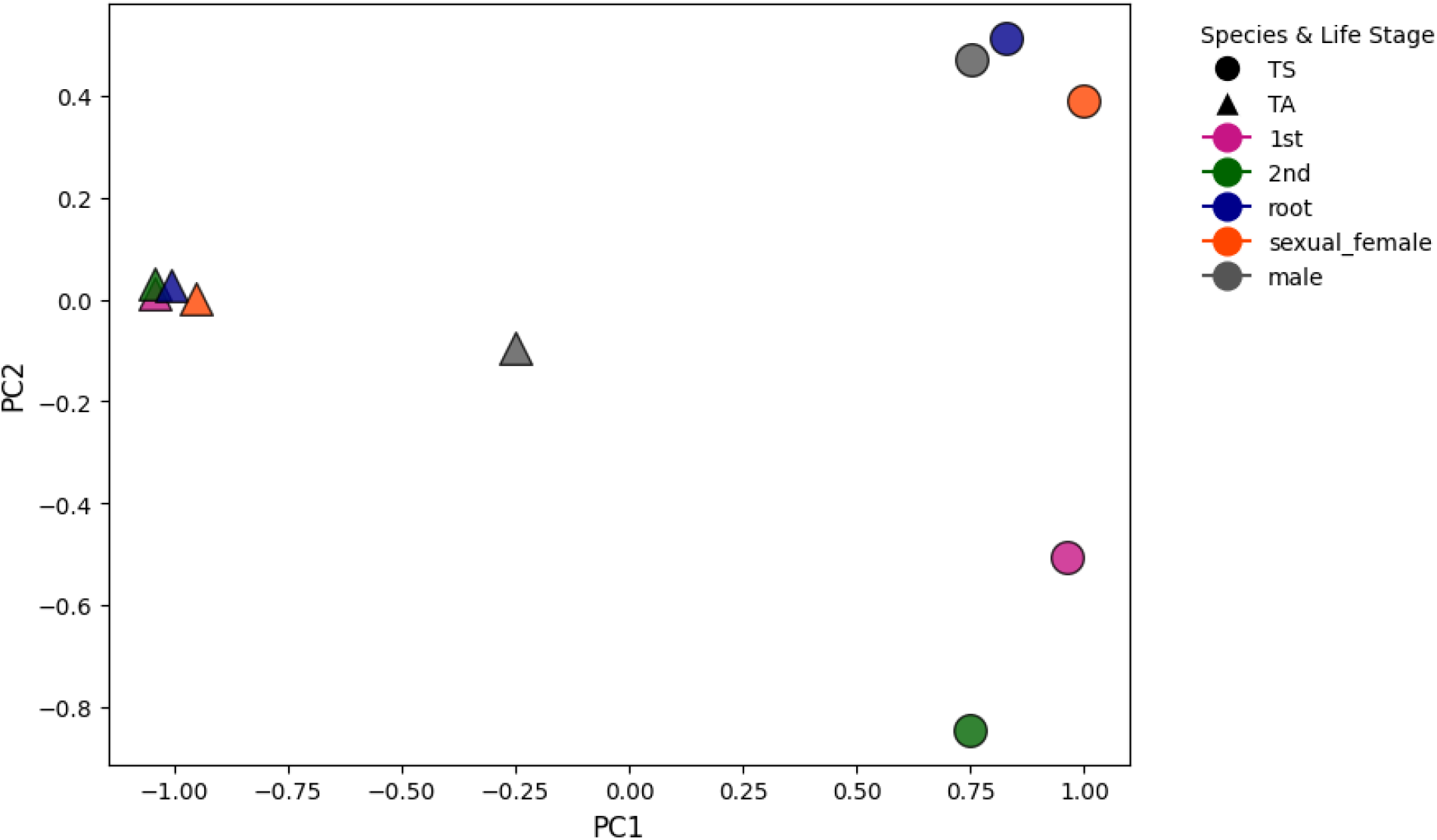
PCoA Analysis of Life Stage Influence on Community Composition. This figure shows the PCoA results for the *Tetraneura* dataset, which includes all life-stage sampling, highlighting the variation in community composition across the different life stages.

### 3.3 *Buchnera* titers in response to life cycle transitions

To quantify *Buchnera* abundance across life stages, we performed qPCR using Ef1a as the reference gene. *Buchnera* titers varied significantly across life stages and species (Figure 4, Kruskal-Wallis H = 23.27, p = 0.0056). Root-generation aphids exhibited higher *Buchnera* titers than gall-generation aphids, with second-generation gall aphids possessing slightly higher titers than first-generation aphids. Sexual females showed lower titers, while males harbored little to no *Buchnera*. However, Dunn’s post hoc test revealed that most pairwise comparisons were not statistically significant (p > 0.05), suggesting overall stability in *Buchnera* titers. An exception was observed in *T. sorini* root aphids, which exhibited significantly different titers compared to *T. akinire* and *T. sorini* males (p = 0.0411 and p = 0.0378, respectively) (Table 2).

**Table 2.** Sample Grouping for Diversity Analysis Based on Aphid Species, Life Stages, and Host Plant. The samples listed below are mixed for 16S rRNA metagenomic sequencing, while the 4th instar and 4-day-old

| ID | Species | Species Group | Life stage | Host group |
| --- | --- | --- | --- | --- |
| S1 | *T. sorini* | TS | 1<sup>st</sup> generation | Elm gall |
| S2 | *T. sorini* | TS | 2nd generation | Elm gall |
| S3 | *T. sorini* | TS | Root generation | Grass root |
| S4 | *T. sorini* | TS | Sexual female | No_feeding |
| S5 | *T. sorini* | TS | Male | No_feeding |
| S6 | *T. akinire* | TA | 1<sup>st</sup> generation | Elm gall |
| S7 | *T. akinire* | TA | 2nd generation | Elm gall |
| S8 | *T. akinire* | TA | Root generation | Grass root |
| S9 | *T. akinire* | TA | Sexual female | No_feeding |
| S10 | *T. akinire* | TA | Male | No_feeding |
|  | *E.* | EH |  |  |
| S11 | *harunire* |  | 2nd generation | Elm gall |
|  | *E.* | EH |  |  |
| S12 | *harunire* |  | Root generation | Grass root |

**Table 3.** Dunn’s Test pairwise comparisons of *Buchnera* Abundance Across Life Stages and Species. The table shows the results of Dunn’s Post Hoc Test for pairwise comparisons of *Buchnera* abundance across different life stages and species. ‘TA’ represents *T. akinire*, ‘TS’ represents *T. sorini*, and the life stages include ‘F’ for Foundress, ‘2G’ for the 2nd Generation, ‘RG’ for Root Generation, ‘M’ for Male, and ‘SF’ for Sexual Female.

|  | TA_F | TS_F | TA-2G | TS-2G | TA-M | TS-M | TA-RG | TS-RG | TA-SF | TS-SF |
| --- | --- | --- | --- | --- | --- | --- | --- | --- | --- | --- |
| TA_F | 1 | 1 | 1 | 1 | 1 | 1 | 1 | 1 | 1 | 1 |
| TS_F | 1 | 1 | 1 | 1 | 1 | 1 | 1 | 0.761235 | 1 | 1 |
| TA-2G | 1 | 1 | 1 | 1 | 1 | 1 | 1 | 1 | 1 | 1 |
| TS-2G | 1 | 1 | 1 | 1 | 0.976025 | 0.917893 | 1 | 1 | 1 | 1 |
| TA-M | 1 | 1 | 1 | 0.976025 | 1 | 1 | 0.145297 | 0.134771 | 1 | 1 |
| TS-M | 1 | 1 | 1 | 0.917893 | 1 | 1 | 0.041077 | 0.037797 | 1 | 1 |
| TA-RG | 1 | 1 | 1 | 1 | 0.145297 | 0.134771 | 1 | 1 | 1 | 1 |
| TS-RG | 1 | 0.761235 | 1 | 1 | 0.041077 | 0.037797 | 1 | 1 | 0.628552 | 1 |
| TA-SF | 1 | 1 | 1 | 1 | 1 | 1 | 1 | 1 | 1 | 1 |
| TS-SF | 1 | 1 | 1 | 1 | 1 | 1 | 1 | 1 | 1 | 1 |

**Figure 4.**
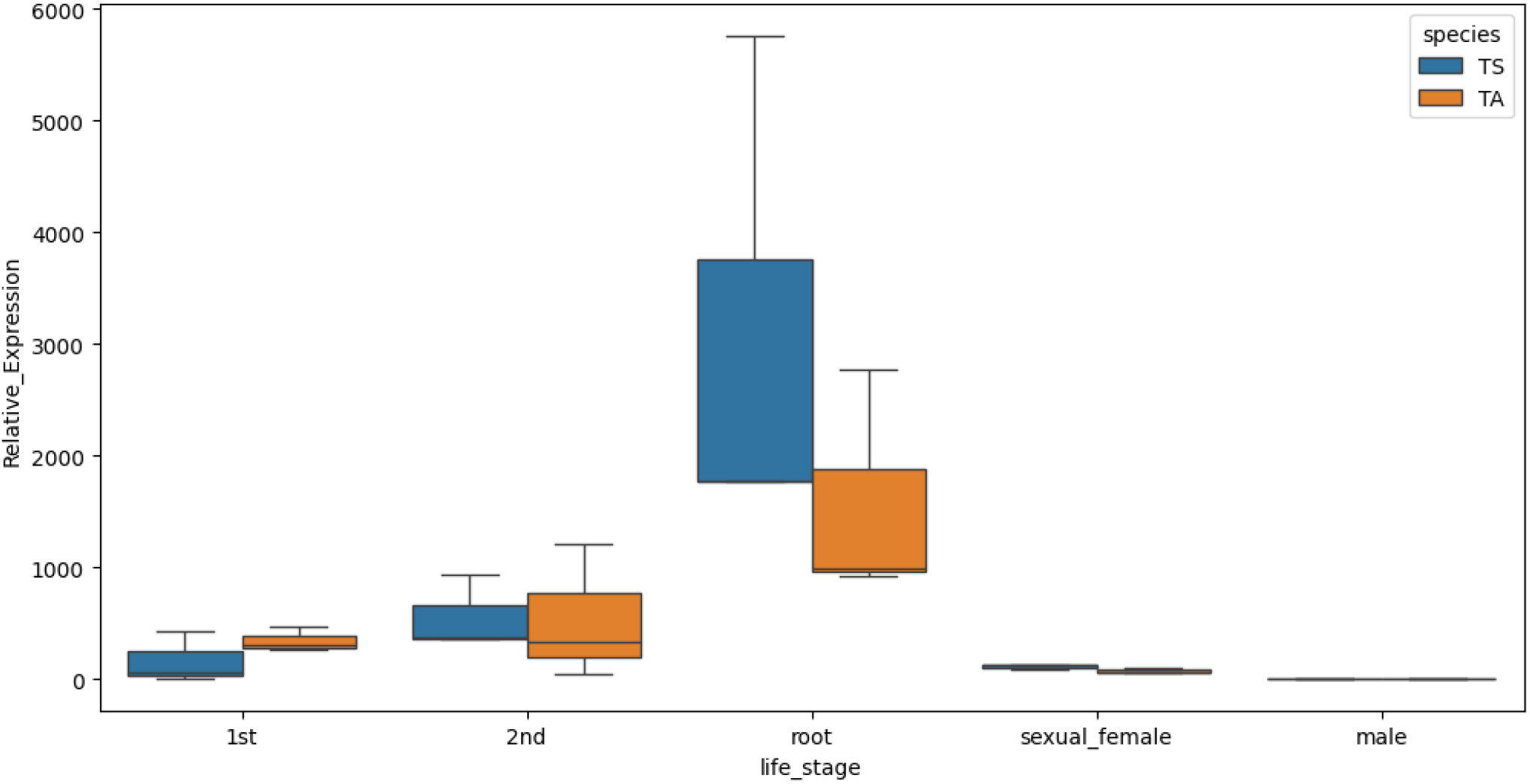
*Buchnera* titers Across Life Stage and Species. *<u>Buchnera</u>* titers of the 4th instar or 4-day-old aphids across the life cycles of *T. sorini* and *T. akinire* were presented and the titers were determined by analyzing the ratio of the single-copy *Buchnera* gene (*Dnak*) to the single-copy aphid gene (ef1a) using the 2^− ΔΔCq^ method for relative expression quantification. The data were analyzed using the Kruskal-Wallis H test, showing that *Buchnera* titers varying significantly across life stages and species (H = 23.27, = 0.0056).

## 2 MATERIALS AND METHODS

### 2.1 Study complex life cycles of gall-forming aphids

*Tetraneura akinire* and *T. sorini* induce closed globular galls on primary host plant, Japanese elm tree, *Ulmus davidiana* var. *japonica*, and seasonally shift host plants between elm trees and Poaceae grasses. In early May in Hokkaido, Japan, a *Tetraneura* foundress hatches from the overwintered egg on the trunk of the elm tree, and each foundress induces a single gall to parthenogenetically produce the second-generation aphids. The gall is well-formed by the foundress before it gives birth to the second-generation aphids. The second-generation aphids grow into winged adults, which migrate through the openings on galls induced by the foundresses, to the roots of Poaceae grasses to produce root-generation aphids. In Hokkaido, the secondary host plants of *T. akinire* and *T. sorini* root generation are *Setaria faberi* and *Miscanthus sinensis* respectively. The root generations reproduce from early summer to autumn, when nymphs develop into winged females (sexuparae). Sexupara is the last generation feeding on the grass roots and the only root-generation aphid that can become winged morph. When the sexupara reaches to the adult stage, it becomes winged adult and migrate back to the Japanese elm and reproduce sexual females and male aphids on the tree trunk. *Eriosoma harunire* shares the same primary host plant, the Japanese elm with *Tetraneura* aphids but induces an open leaf-curl gall to produce the second-generation and the third-generation aphids in the gall. The secondary host plant of *E. harunire* is *Plantago asiatica*.

### 2.2 sample collection

Galls including the phases of foundresses and the second-generation aphids were accordingly collected from the elm tree as the galling generation, and apterous exule aphids and sexupara nymphs were collected on the grass roots as the root generation. However, since the instars of sexual-generation aphids were indecipherable, we first collected the tree barks with newborn sexual nymphs on the day of the first peak of sexupara reproduction on elm tree, and sexual nymphs were collected from the tree barks three days later to avoid aged samples. Nonetheless, due to the low population of *E. harunire*, sexual generations of *E. harunire* were not included. Here only the 4^th^ instar of aphids in both galling and root generation were used for *Buchnera* titer quantification, together with 4-day old sexual nymphs which are presumably around 4^th^ instar. All samples were maintained in 99% ethanol at −20 degrees Celsius for later analysis, and details of sample collection were listed in Table 1.

### 2.3 DNA extraction, library preparation, and sequencing

#### 16S rDNA amplicon sequencing

| The genomic DNA (gDNA) of aphid samples were extracted respectively by different life stages and instars of aphids using QIAGEN DNeasy Blood & Tissue Kit following the manufacturer’s protocol. The target 16S rDNA amplicons were amplified using the standard primers for V1-V2, and V3-V4 regions in Zymo Research Quick-16STM NGS Library Prep Kit. The PCR annealing step was slightly modified to 30 cycles due to low input DNA concentration. TapeStation HS D5000 (Agilent Technologies) was used to screen quality of the DNA amplicons. From the step of DNA cleanup, library preparation was performed using TruSeq Nano DNA Prep following the instruction of library preparation for DNA amplicons. Before normalizing and pooling libraries, TapeStation was employed to screen the library quality. The libraries were sequenced using the Illumina MiSeq platform at Trans-Omics Facility, National Institute for Basic Biology, Japan.

### 2.4 Bioinformatic analysis

#### Analysis of 16S rDNA amplicon sequencing

The raw paired-end reads of 16S rDNA amplicon sequencing were demultiplexed according to their unique barcode sequences. As V1-V2 and V3-V4 reads were pooled and sequenced, the respective primer sequences were used to extract each pooling: 16S V1-V2 Primer Sequences (adapters not included) –27f (AGRGTTYGATYMTGGCTCAG, 20 bp) and 341r (CTGCWGCCHCCCGTAGG, 17 bp); 16S V3-V4 Primer Sequences (adapters not included) – 341f(CCTACGGGDGGCWGCAG, CCTAYGGGGYGCWGCAG, 17bp) and 806r (GACTACNVGGGTMTCTAATCC, 24 bp). The forward primer 341f is a mixture of the two sequences listed according to the kit manual. The quality of the reads was checked by FastQC and previewed by MAFFT. Paired-end reads were imported to Qiime2 (Quantitative Insights Into Microbial Ecology). However, due to quality of the reads, single-end reads were selected. DADA2 plugin (Callahan *et al*. 2016) was assigned to denoise samples by trimming the sequences by 17 bp from 5’ end of reverse reads with truncation parameters of --p-trunc-len 200 as well as removing chimeras. The quality-filtered reads were summarized to generate a feature table with representative sequences ranging from 211bp to 401bp. Alpha and Beta diversities were 10alculated by the q2-diversity plugin. The feature-classifier plugin was applied to conduct taxonomic assignment of representative sequences against the SILVA database (silva-138.2-99) and the taxonomic profiles of each sample were visualized using the qiime taxa barplot command. The codes for analysis were deposited in the Github workflow (https://tinyurl.com/59eams59).

The 16S rRNA gene sequences (V1-V2 and V3-V4 regions) of symbiotic bacteria from *Tetraneura sorini, Tetraneura akinire*, and *Eriosoma harunire* aphids have been deposited in the NCBI Sequence Read Archive (SRA) under accession number PRJNA1232350.

### 2.5 qPCR and statistical analyses

The genomic DNA (gDNA) from aphid samples was extracted using the QIAGEN DNeasy Blood & Tissue Kit. Quantification of *Buchnera* titer was performed using SYBR Green-based qPCR, targeting the single-copy nuclear genes dnaK in *Buchnera* and elongation factor-1 alpha (EF1α) in aphids. Primers targeting *Buchnera* dnaK were designed based on *Buchnera* genome sequencing data (Tong *et al*. 2024): buch_dnaK_erio_f (5’-ACA GCT ACG GCT TCA TCT GG-3’) and buch_dnaK_erio_r (5’-GTA GGT GGT CAA ACA AGA ATG CC-3’). Primers targeting reference aphid *EF1a* gene were designed based unpublished amplicon sequencing data: ef1a172-erio-f (5’-ACT GAA CCA CCA TAC AGT GAA GT-3’) and ef1a172-erio-r (5’-TCC CTT GAA CCA TGG CAT TT-3’). The qPCR was performed in technical duplicates using the Thunderbird SYBR qPCR Mix (Toyobo) and Mx399P QPCR system (Agilent, Santa Clara, USA).

To calculate the *Buchnera* titer, the Cq values of the target *Buchenra* single-copy gene Dnak and reference aphid single-copy gene Ef1a were processed, replacing “No Ct” with NaN. ΔCq was calculated as the difference between Cq_dnak and Cq_ef1a, with missing values substituted by a maximum Cq value of 40. A reference ΔCq was derived as the mean ΔCq of the 1st life stage for the species “TS.” ΔΔCq was then calculated by subtracting this reference value from the individual ΔCq values, and relative expression was computed as 2^(−ΔΔCq), with missing Cq_dnak values set to 0. Normality was checked using the Shapiro-Wilk test. Since the data did not meet normality assumptions, a Kruskal-Wallis H test was applied. Dunn’s post-hoc test was used for pairwise comparisons with Bonferroni correction, and visualizations, including boxplots, were created to examine the distributions of relative expression across different life stages and species. The statistical analysis was computed on Google Colab, and the codes and data are open listed (https://tinyurl.com/4jernnw5).

## 4 Discussion

Our findings highlight significant differences in microbial community among three gall-forming aphid species, *T. akinire, T. sorini*, and *E. harunire*, across various life cycle stages and host plants. While *Buchnera* remains the dominant symbiont in most samples, we observed a striking shift in microbial composition in male aphids, particularly in *T. akinire*, where *Regiella* becomes the dominant bacterium. In contrast to Xu et al. (2020), our study observed a shift towards *Regiella* dominance in male aphids, suggesting that microbial dynamics are more influenced by aphid life stages and reproductive strategies than by host plant effects. This discrepancy could be attributed to variations in sample types, geographical locations, or research methodologies. The decrease in the relative abundance of *Buchnera* in *Tetraneura* males aligns with previous studies in *Pseudoregma bambucicola*, where male aphids exhibited increased bacterial diversity and a reduced abundance of *Buchnera* (Liu et al., 2022). The presence of *Regiella* in male aphids, in contrast to the dominance of *Buchnera* in other life stages, suggests that microbial community compositions are tightly regulated by the aphid’s complex life cycle. These shifts likely reflect the nutritional and ecological demands of specific life stages, such as the male sexual stage, which may be linked to changes in aphid metabolism or reproductive needs. The fact that males do not directly produce offspring may be related to the decreased relative abundance of *Buchnera*. Interestingly, despite these microbial shifts, we found no significant differences in microbial diversity across life stages (e.g., gall generation vs. root generation). This suggests that, although aphids shift between different host plants, their microbial communities remain relatively stable. This stability may reflect a well-integrated host-symbiont system, where *Buchnera* is maternally transmitted and continues to provide essential nutrients, even though other bacteria like *Regiella* become more prominent during certain life stages. *Buchnera* titers in *Tetraneura* exhibited significant variability across different species and life stages, with root-generation aphids showing higher titers compared to gall-generation aphids. The lower *Buchnera* abundance in gall-generation aphids, despite the availability of substantial nutritional resources in the galls (Koyama et al. 2004; Suzuki et al. 2009), suggests that the nutritional quality of galls may not be as high as previously assumed, leading to a lower expression of *Buchnera* during these stages. Contrary to the hypothesis proposed by Suzuki et al. (2009), which suggests that galls provide superior nutritional conditions compared with healthy leaves, our findings show that gall-generation aphids do not exhibit higher *Buchnera* titers. This suggests that the nutritional conditions in galls may not be high in terms of absolute titers of amino acids, particularly when we observe that second-generation aphids, which developed by feeding on galls, did not show elevated *Buchnera* titers compared to the generations feeding on grass roots. Additionally, the marked reduction or near-absence of *Buchnera* in male aphids further underscores how microbial dynamics are closely tied to host biology. These further challenges the idea that galls are a better nutritional resource compared with the secondary host plants. Understanding the ecological and evolutionary drivers of aphid-microbe interactions will provide deeper insights into host-microbe co-evolution and the adaptive significance of microbial diversity in aphids. Future studies should aim to elucidate the functional implications of these microbial shifts, particularly regarding the role of *Regiella* and other secondary symbionts in male aphids.

## ACKNOWLEDGEMENTS

X.T. is supported by the Special Postdoctoral Researcher Fellowship from Institute of Physical and Chemical Research (RIEKN) in Japan and JSPS Start-up Research Funding (Project Number: 22K20588). Computational resources were provided by the Data Integration and Analysis Facility, National Institute for Basic Biology, and the Research Center for Computational Science, Okazaki, Japan (Project: NIBB, 24-IMS-C279). The authors thank Dr. Nishide Hiroyo at the National Institute for Basic Biology for her support with the computational platform during the project.

## Author contributions

X.T, S.A and S.S designed the research; X.T collected the samples with further identification by S.A; X.T performed data analysis; X.T wrote the paper with contribution from S.A, T.N and M.A.

## Data availability

The datasets generated and analyzed during the current study are available in the NCBI GenBank repository, under the BioProject accession number PRJNA1232350.

## REFERENCES

Stewart, F. J., Newton, I. L., & Cavanaugh, C. M. (2005). Chemosynthetic endosymbioses: Adaptations to oxic-anoxic interfaces. Trends in Microbiology, 13(9), 439–448. 10.1016/j.tim.2005.07.007

Relman, D. (2008). ‘Til death do us part’: Coming to terms with symbiotic relationships. Nature Reviews Microbiology, 6, 721–724. 10.1038/nrmicro1990

Forcada, J., Trathan, P. N., & Murphy, E. J. (2008). Life history buffering in Antarctic mammals and birds against changing patterns of climate and environmental variation. Global Change Biology, 14, 2473–2488. 10.1111/j.1365-2486.2008.01678.x

Bell, G. (2010). Fluctuating selection: The perpetual renewal of adaptation in variable environments. Philosophical Transactions of the Royal Society B: Biological Sciences, 365, 87–97. 10.1098/rstb.2009.0150

Nettle, D., & Bateson, M. (2015). Adaptive developmental plasticity: What is it, how can we recognize it and when can it evolve? Proceedings of the Royal Society B, 282, 20151005. 10.1098/rspb.2015.1005

Wilbur, H. M. (1980). Complex life cycles. Annual Review of Ecology and Systematics, 11, 67–93. 10.2307/2096903

Moran, N. A. (1992). The evolutionary maintenance of alternative phenotypes. The American Naturalist, 139(5), 971–989. 10.1086/285369

Minelli, A., & Fusco, G. (2010). Developmental plasticity and the evolution of animal complex life cycles. Philosophical Transactions of the Royal Society B: Biological Sciences, 365, 631–640. 10.1098/rstb.2009.0268

Auld, S., & Tinsley, M. (2015). The evolutionary ecology of complex lifecycle parasites: Linking phenomena with mechanisms. Heredity, 114, 125–132. 10.1038/hdy.2014.84

Rauch, G., Kalbe, M., & Reusch, T. B. H. (2005). How a complex life cycle can improve a parasite’s sex life. Journal of Evolutionary Biology, 18(4), 1069–1075. 10.1111/j.1420-9101.2005.00895.x

Parker, B. J., et al. (2019). A laterally transferred viral gene modifies aphid wing plasticity. Current Biology, 29(12), 2098–2103.e5. 10.1016/j.cub.2019.05.070

Benesh, D. P., Chubb, J. C., Lafferty, K. D., & Parker, G. A. (2022). Complex life-cycles in trophically transmitted helminths: Do the benefits of increased growth and transmission outweigh generalism and complexity costs? Current Research in Parasitology & Vector-Borne Diseases, 2, 100085. 10.1016/j.crpvbd.2022.100085

Hanna, L., & Abouheif, E. (2021). The origin of wing polyphenism in ants: An eco-evo-devo perspective. In S. F. Gilbert (Ed.), Current Topics in Developmental Biology (Vol. 141, pp. 279–336). Academic Press. 10.1016/bs.ctdb.2020.12.004

Moran, N. A. (1994). Adaptation and constraint in the complex life cycles of animals. Annual Review of Ecology and Systematics, 25, 573–600. 10.2307/2097325

Gandon, S. (2004). Evolution of multihost parasites. Evolution, 58(3), 455–469. 10.1111/j.0014-3820.2004.tb01669.x

Bonett, R. M., & Blair, A. L. (2017). Evidence for complex life cycle constraints on salamander body form diversification. Proceedings of the National Academy of Sciences, 114(37), 9936–9941. 10.1073/pnas.1703877114

Jousselin, E., Genson, G., & Coeur d’Acier, A. (2010). Evolutionary lability of a complex life cycle in the aphid genus Brachycaudus. BMC Evolutionary Biology, 10, 295. 10.1186/1471-2148-10-295

Favret, C. (2022). Aphid species file (Version 5.0/5.0). Retrieved from http://aphid.speciesfile.org

Akimoto, S. I. (1985). Taxonomic study on gall aphids, Colopha, Paracolopha and Kaltenbachiella (Aphidoidea: Pemphigidae) in East Asia, with special reference to their origins and distributional patterns. Insecta Matsumurana, 31, 1–79.

Moran, N. A. (1988). The evolution of host-plant alternation in aphids: Evidence for specialization as a dead end. The American Naturalist, 132(5), 681–706. 10.1086/284881

Stern, D. L. (2008). Aphids. Current Biology, 18(R504–R505). 10.1016/j.cub.2008.03.052

Aoki, S., & Kurosu, U. (2010). A review of the biology of Cerataphidini (Hemiptera, Aphididae, Hormaphidinae), focusing mainly on their life cycles, gall formation, and soldiers. Psyche: A Journal of Entomology, 2010, 380351. 10.1155/2010/380351

Ward, S. A., Leather, S. R., Pickup, J., & Harrington, R. (1998). Mortality during dispersal and the cost of host-specificity in parasites: How many aphids find hosts? Journal of Animal Ecology, 67(4), 763–773. 10.1046/j.1365-2656.1998.00238.x

Stone, G. N., Schönrogge, K., Atkinson, R. J., Bellido, D., & Pujade-Villar, J. (2002). The adaptive significance of insect gall morphology. Trends in Ecology & Evolution, 18(10), 512–522. 10.1016/S0169-5347(03)00137-3

Harris, M. O., & Pitzschke, A. (2020). Plants make galls to accommodate foreigners: Some are friends, most are foes. New Phytologist, 225(4), 1852–1872. 10.1111/nph.16340

Korgaonkar, A., Han, C., Lemke, A., et al. (2021). A novel family of secreted insect proteins linked to plant gall development. Current Biology, 31(9), 1836–1849.e12. 10.1016/j.cub.2021.03.021

Hille Ris Lambers, D. (1967). Notes on California aphids, with descriptions of new genera and new species (Homoptera: Aphididae). Hilgardia, 37(15), 569–623.

Heie, O. E. (1997). The evolutionary history of aphids and a hypothesis on the coevolution of aphids and plants. Bollettino di Zoologia Agraria e di Bachicoltura, 28(2), 149–155.

Suzuki, D. K., Fukushi, Y., & Akimoto, S. I. (2009). Do aphid galls provide good nutrients for the aphids?: Comparisons of amino acid concentrations in galls among Tetraneura species (Aphididae: Eriosomatinae). Arthropod-Plant Interactions, 3, 241–247. 10.1007/s11829-009-9064-9

Koyama, Y., Yao, I., & Akimoto, S. I. (2004). Aphid galls accumulate high concentrations of amino acids: A support for the nutrition hypothesis for gall formation. Entomologia Experimentalis et Applicata, 113(1), 35–44. 10.1111/j.0013-8703.2004.00207.x

Williams, A. G., & Whitham, T. G. (1986). Premature leaf abscission: An induced plant defense against gall aphids. Ecology, 67(6), 1619–1627. 10.2307/1939093

Goggin, F. L. (2007). Plant–aphid interactions: Molecular and ecological perspectives. Current Opinion in Plant Biology, 10(4), 399–408. 10.1016/j.pbi.2007.06.004

Douglas, A. E. (1998). Nutritional interactions in insect-microbial symbioses: Aphids and their symbiotic bacteria Buchnera. Annual Review of Entomology, 43, 17–37.

Douglas, A. E., François, C. L. M. J., & Minto, L. B. (2006). Facultative ‘secondary’ bacterial symbionts and the nutrition of the pea aphid, Acyrthosiphon pisum. Physiological Entomology, 31(3), 262–269. 10.1111/j.1365-3032.2006.00516.x

Shigenobu, S., Watanabe, H., Hattori, M., et al. (2000). Genome sequence of the endocellular bacterial symbiont of aphids Buchnera sp. APS. Nature, 407, 81–86. 10.1038/35024074

Manzano□Marín, A., Szabó, G., Simon, J. C., Horn, M., & Latorre, A. (2017). Happens in the best of subfamilies: Establishment and repeated replacements of coLobligate secondary endosymbionts within Lachninae aphids. Environmental Microbiology, 19(1), 393–408.

Manzano-Marín, A., et al. (2023). Co-obligate symbioses have repeatedly evolved across aphids, but partner identity and nutritional contributions vary across lineages. Peer Community Journal, 3.

McLean, A. H. C., et al. (2017). Cascading effects of herbivore protective symbionts on hyperparasitoids. Ecological Entomology, 42(5), 601–609.

Douglas, A. E., Minto, L. B., & Wilkinson, T. L. (2001). Quantifying nutrient production by the microbial symbionts in an aphid. Journal of Experimental Biology, 204(2), 349–358.

Koga, R., Tsuchida, T., & Fukatsu, T. (2003). Changing partners in an obligate symbiosis: A facultative endosymbiont can compensate for loss of the essential endosymbiont Buchnera in an aphid. Proceedings of the Royal Society B: Biological Sciences, 270(1533), 2543–2550.

Hansen, A. K., & Moran, N. A. (2011). Aphid genome expression reveals host–symbiont cooperation in the production of amino acids. Proceedings of the National Academy of Sciences, 108(7), 2849–2854.

Wilson, A. C., et al. (2010). Genomic insight into the amino acid relations of the pea aphid, Acyrthosiphon pisum, with its symbiotic bacterium Buchnera aphidicola. Insect Molecular Biology, 19, 249–258.

Francis, F., Haubruge, E., Hastir, P., & Gaspar, C. (2001). Effect of aphid host plant on development and reproduction of the third trophic level, the predator Adalia bipunctata (Coleoptera: Coccinellidae). Environmental Entomology, 30(5), 947–952.

Zhang, Y. C., Cao, W. J., Zhong, L. R., Godfray, H. C. J., & Liu, X. D. (2016). Host plant determines the population size of an obligate symbiont (Buchnera aphidicola) in aphids. Applied and Environmental Microbiology, 82(8), 2336–2346.

Parven, N., Akimoto, S. I., Yao, I., & Kanbe, T. (2024). Effects of climate change and high temperature heat stress on aphid fitness traits and symbionts.

Oliver, K. M., Russell, J. A., Moran, N. A., & Hunter, M. S. (2003). Facultative bacterial symbionts in aphids confer resistance to parasitic wasps. Proceedings of the National Academy of Sciences, 100(4), 1803–1807.

Vorburger, C., Sandrock, C., Gouskov, A., Castaneda, L. E., & Ferrari, J. (2009). Genotypic variation and the role of defensive endosymbionts in an all-parthenogenetic host–parasitoid interaction. Evolution, 63(6), 1439–1450.

Yorimoto, S., Hattori, M., Kondo, M., & Shigenobu, S. (2022). Complex host/symbiont integration of a multi-partner symbiotic system in the eusocial aphid Ceratovacuna japonica. iScience, 25(12).

Liu, S., Liu, X. B., Zhang, T. T., Bai, S. X., He, K. L., Zhang, Y. J., Francis, F., & Wang, Z. Y. (2024). Effects of host plants on aphid feeding behavior, fitness, and Buchnera aphidicola titer. Insect Science.

Stewart, C. J., et al. (2018). Temporal development of the gut microbiome in early childhood from the TEDDY study. Nature, 562(7728), 583–588.

Hammer, T. J., EastonLCalabria, A., & Moran, N. A. (2023). Microbiome assembly and maintenance across the lifespan of bumble bee workers. Molecular Ecology, 32(3), 724–740.

Obadia, B., et al. (2017). Probabilistic invasion underlies natural gut microbiome stability. Current Biology, 27(13), 1999–2006.

Vega, N. M., & Gore, J. (2017). Stochastic assembly produces heterogeneous communities in the Caenorhabditis elegans intestine. PLOS Biology, 15(3), e2000633.

Song, S. J., et al. (2013). Cohabiting family members share microbiota with one another and with their dogs. eLife, 2, e00458.

Kwong, W. K., & Moran, N. A. (2016). Gut microbial communities of social bees. Nature Reviews Microbiology, 14(6), 374–384.

Hammer, T. J., De Clerck-Floate, R., Tooker, J. F., Price, P. W., Miller, D. G., & Connor, E. F. (2021). Are bacterial symbionts associated with gall induction in insects? Arthropod-Plant Interactions, 15, 1–12.

Xu, T. T., Jiang, L. Y., Chen, J., & Qiao, G. X. (2020). Host plants influence the symbiont diversity of Eriosomatinae (Hemiptera: Aphididae). Insects, 11(4), 217.

